# Gsα stimulation of mammalian adenylate cyclases regulated by their hexahelical membrane anchors

**DOI:** 10.1101/694877

**Authors:** Anubha Seth, Manuel Finkbeiner, Julia Grischin, Joachim E. Schultz

## Abstract

Mammalian adenylate cyclases (ACs) are pseudoheterodimers with dissimilar hexahelical membrane-anchors, isoform-specifically conserved for more than half a billion years. We exchanged both membrane anchors of the AC isoform 2 by the isosteric quorum-sensing receptor from *Vibrio*, CqsS, which has a ligand, *Cholera*-Autoinducer-1 (CAI-1). In the chimera, AC activity was stimulated by Gsα, CAI-1 had no effect. Surprisingly, CAI-1 inhibited Gsα stimulation. We report that Gsα stimulation of human AC isoforms 2, 3, 5, and 9 expressed in Sf9 cells is inhibited by serum as is AC activity in membranes from rat brain cortex. AC2 activation by forskolin or forskolin/Gsα was similarly inhibited. Obviously, serum contains as yet unidentified factors affecting AC activity. The data establish a linkage in ACs, in which the membrane anchors, as receptors, transduce extracellular signals to the cytosolic catalytic dimer. A mechanistic three state model of AC regulation is presented compatible with all known regulatory inputs into mammalian ACs. The data allow designating the membrane anchors of mammalian ACs as orphan receptors and establish a new level of AC regulation.

## Introduction

The first amino acid sequence of a mammalian adenylate cyclase identified two similar catalytic domains (C1 and C2) and two dissimilar hexahelical membrane anchors (TM1 and TM2) which were proposed to possess a channel or transporter-like function, properties, which were never confirmed (Krupinski, Coussen et al., 1989). Subsequently, nine genes for mAC isoforms were identified, indicating substantial subfunctionalization during evolution (mAC isoforms 1 - 9; (Dessauer, Watts et al., 2017, Sinha & Sprang, 2006)). mACs are the cellular effector proteins for many hormones that signal via G-protein-coupled receptors and their regulation has received broad attention (Dessauer et al., 2017, Sinha & Sprang, 2006). The catalytic center of mACs is formed at the C1/C2 dimer interface. Most biochemical studies have used the startling observation that the separately expressed C1/C2 catalytic domains are regulated by Gsα, i.e. the membrane anchors appear dispensable for catalysis and regulation (Tang & Gilman, 1995). Why then 2 x 6 transmembrane spans, when 1 or 2 would have been sufficient for membrane-anchoring? The evolutionary conservation of the membrane domains for more than half a billion years justifies searching for a physiological function beyond membrane-anchoring (Bassler, Schultz et al., 2018, Beltz, Bassler et al., 2016, Ziegler, Bassler et al., 2017).

Recently, we replaced the 6TM domain of the mycobacterial Rv1625c AC, a monomeric progenitor of mACs, by the hexahelical quorum-sensing receptor CqsS from *Vibrio*, generating a CqsS-AC Rv1625c chimera (here and in the following, CqsS is used to denote the ligand-binding membrane domain of CqsS, not the full receptor protein (Beltz et al., 2016, Ziegler et al., 2017)). *The ligand for CqsS, Cholera*-Auto-Inducer-1 [CAI-1; (S)-3-hydroxy-tridecan-4-one], stimulated AC activity in the chimera (Beltz et al., 2016). Subsequently, we characterized a family of conserved cyclase-transducing-elements (CTEs) which are indispensable for signal transduction (Ziegler et al., 2017). They are isoform-specifically conserved in mACs, supporting the notion that the AC membrane domains may be ligand receptors (Bassler et al., 2018, Beltz et al., 2016, Ziegler et al., 2017). Here, we asked whether AC regulation by CqsS is maintained in a chimera, in which the TM1 and TM2 domains in human AC2 (hAC2) are replaced by CqsS, generating CqsS-hAC2. We report that stimulation by Gsα is preserved whereas the ligand CAI-1 does not, by itself, affect basal activity. Surprisingly, CAI-1 inhibits Gsα stimulation in CqsS-hAC2. We further show that the Gsα stimulation of the hAC2 holoenzyme is similarly inhibited by human serum. Serum also inhibited Gsα activation of hAC isoforms 3, 5, and 9, and AC activity in a rat brain cortical membrane indicating that the AC membrane domains are orphan ligand receptors.

## Results

### CqsS serves as a ligand receptor for hAC2

We generated a pseudoheterodimeric chimera of hAC2 in which both 6TMs were isosterically replaced by the 6TM quorum-sensing receptor CqsS from *Vibrio* (CqsS-hAC2). The point of transition between CqsS and hAC2 was at the respective hAC2 CTEs in front of the catalytic domains C1a and C2, thus maintaining all structural features potentially required for signaling (Fig. 1 left; (Beltz et al., 2016, Qi, Sorrentino et al., 2019, Ziegler et al., 2017)). Two questions were obvious: a) Is such a chimera expressed in *Escherichia coli* although bacterial expression of mammalian ACs has so far proven impossible, and b) Is regulation by Gsα and the quorum-sensing ligand CAI-1 maintained? The chimera was expressed in *E. coli* and had robust basal activity indicating that native mAC-like features were maintained (Fig. 1). With Mn^2+^ as divalent cation basal AC activity was stimulated by constitutively active Gsα (Q227L, below termed Gsα (Graziano, Freissmuth et al., 1989)) demonstrating formation of a productive C1/C2 catalytic dimer. A concentration-response curve showed a 2.3-fold increase in activity with an EC_50_ of around 200 nM Gsα (Fig. 1). AC2 expressed in Sf9 cells is stimulated by Gsα about 5 to 12-fold (Tang, Stanzel et al., 1995, Weitmann, Schultz et al., 2001). The quantitatively differing responses compared to AC2 may be due to the replacement of the dissimilar TM1 and TM2 domains by two identical CqsS receptors. The quorum-sensor ligand CAI-1, up to 100 μM, failed to affect AC activity. This posed two questions: a) Are the catalytic domains of mACs at all capable of operating as output domains for transmembrane signals? b) Which biochemical differences between bacterial and mammalian ACs exist, which might explain the divergent results obtained with the CqsS-Rv1625c AC chimera (Beltz et al., 2016)?

**Figure 1.**
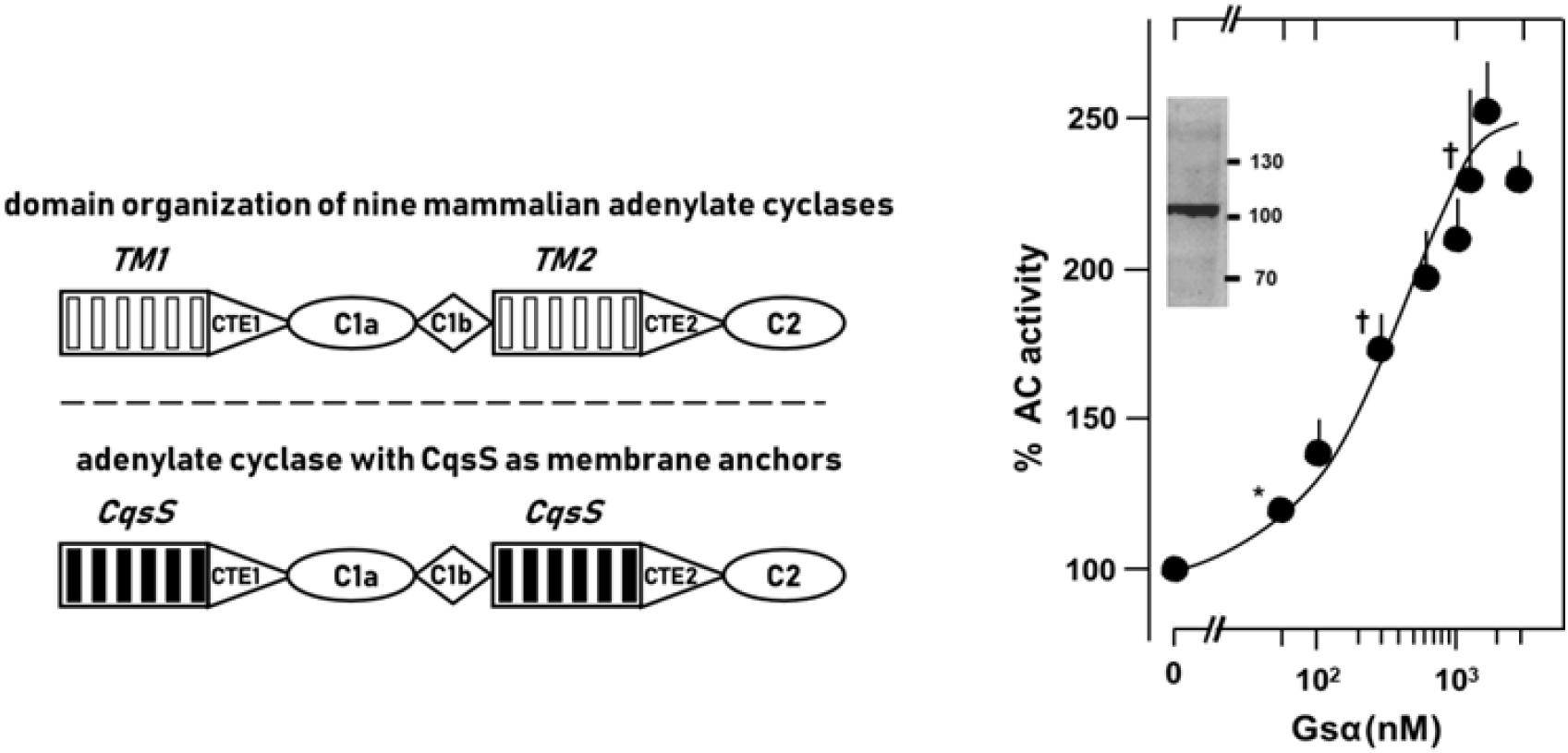
Stimulation of the chimera CqsS-hAC2 by Gsα. **Left**: general domain organization of the membrane-delimited adenylate cyclases (top) and scheme of the chimera between CqsS as membrane anchors and hAC2 (bottom). The 6TM regions are idealized, CTE1 and CTE2 denote the cyclase-transducing-elements, indispensable for signal transduction (Beltz et al., 2016, Ziegler et al., 2017), C1a and C2 are the respective catalytic domains, C1b connects C1a to TM2 (construct boundaries: MRGSHis_6_-GS–CqsS (F166L)_1-181_-hAC2_221-603_-RS-CqsS (F166L)_1-181_-hAC2_836-1091_). **Right:** Gsα concentration-response curve. Basal activity (100 %) was 27.2 ± 2.6 pmol cAMP·mg^−1^min^1^. Half-maximal stimulation was around 147.6 nM Gsα. Error bars denote S.E.M. of 2-7 separate experiments (expressions). Significances: ‡: p < 0.001 compared to basal. For clarity, not all significances are marked. Insert: Western blot of the chimera with an anti-His6-antibody indicating absence of proteolysis. MW standards are at right.

A crucial difference between bacterial and mammalian ACs are the dissociation constants of the catalytic domains. Bacterial catalytic domains usually have a high ‘self’-affinity (Kd ≤10^−7^ M) and are active when conformationally unconstrained (Bassler et al., 2018, Beltz et al., 2016, Guo, Seebacher et al., 2001, Ziegler et al., 2017). Similarly, the individual C1 and C2 domains of mACs have a high propensity for self-association, i.e. C1 preferentially associates with C1 and C2 with C2, as documented in the first mAC crystal structure, a C2 homodimer (Zhang, Liu et al., 1997). However, homodimers of C1 or C2 are inactive (Yan, Hahn et al., 1996). The actual affinity between the C1 and C2 catalytic domains in mACs is rather low (Kd ≤10^−5^ M (Dessauer, Chen-Goodspeed et al., 2002, Hatley, Benton et al., 2000, Tesmer, Sunahara et al., 1997). AC stimulation by Gsα is tantamount to an increase in the apparent affinity of C1 and C2 for each other by approximately two orders of magnitude (Dessauer et al., 2002, Hatley et al., 2000, Ritt & Sivaramakrishnan, 2016, Tesmer et al., 1997). Therefore, a provocative interpretation for the lack of a CAI-1 effect would be that CqsS receptor activation causes conformational changes which interfere with Gsα stimulation. This was tested next.

### The CAI-1 receptor regulates stimulation of hAC2 by Gsα

In presence of 5 or 10 μM CAI-1, Gsα activation of CqsS-hAC2 was significantly attenuated (Fig. 2A). Inhibition was instantaneous. The effect was ligand-specific (Beltz et al., 2016) and reversible (Fig. 2B). In presence of 5 or 10 μM CAI-1, the EC_50_ concentrations for Gsα were increased 1.5- and 4-fold, respectively (Fig. 2A). Concomitantly, CAI-1 at 10 μM diminished the maximal Gsα response by 20% (Fig. 2A). The action of CAI-1 was reversible, as determined by re-assaying membranes which were stimulated, washed and re-isolated (Fig. 2B). The data supported the hypothesis that CqsS receptor activation interfered with activation by Gsα.

**Figure 2.**
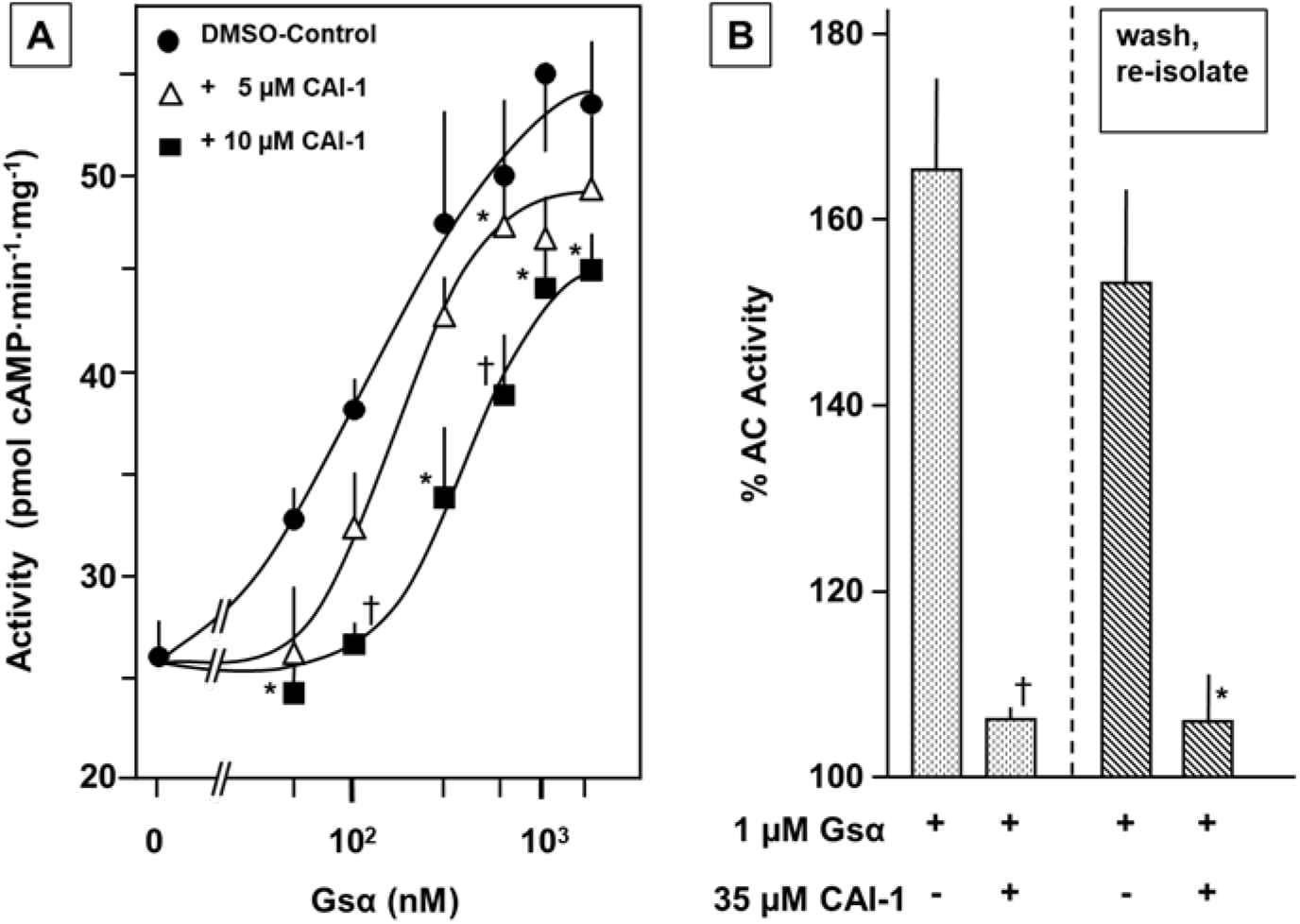
**A)** Effect of CAI-1 on the stimulation of the chimera CqsS-hAC2 by Gsα. Filled circles, Gsα alone; open triangles, + 5 μM CAI-1; filled squares, + 10 μM CAI-1. Error bars denote S.E.M. of 3-4 separate experiments (expressions). Significances: *, p <0.05; †, p < 0. 01 compared to the respective Gsα stimulation. **B)** Inhibition of the Gsα response by CAI-1 is reversible. After stimulation of CqsS-hAC2 by 1 μM Gsα ± 10 μM CAI-1 for 15 min (left) membranes were washed and re-isolated by ultracentrifugation (total time required about 150 min). Re-stimulation was for 15 min. Basal activity (100%) corresponded to 48.2 ± 9.1 (primary stimulation) and 32.3 ± 5.1 pmol cAMP·mg^−1^·min^−1^ (washed membranes). Error bars denote S.E.M. of 5 separate experiments (expressions). Significances: *, p <0.05; †, p < 0.01 compared to the respective Gsα stimulation.

Next, a concentration-response curve for CAI-1 in the presence of 1 μM Gsα was carried out. At 35 μM CAI-1, activation of the CqsS-hAC2 chimera by Gsα was almost abrogated (Fig. 3A). The IC_50_ for CAI-1 inhibition was 6.3 μM, i.e. about 15-fold higher than for the stimulatory effect of CAI-1 in the CqsS-Rv1625c AC (Beltz et al., 2016). This might be due to the fact that we went from a homodimeric CqsS-Rv1625c AC to a linked pseudoheterodimeric CqsS-hAC2 chimera. To exclude that CAI-1 might have obstructed the formation of the catalytic dimer or its interaction with Gsα, the effect of CAI-1 on Gsα stimulation of native hAC2 expressed in Sf9 cells was tested. CAI-1 neither affected basal nor Gsα-stimulated AC activity (Fig. 3B). This unequivocally demonstrated that a) CAI-1 did not interfere in formation of the catalytic dimer in the CqsS-hAC2 chimera, b) CAI-1 did not interact with Gsα and impair its function, and c) CAI-1 did not impair the interactions between the catalytic dimer and Gsα. From these data we can conclude that the effect of CAI-1 in the CqsS-hAC2 chimera was mediated via the CqsS-receptor.

**Figure 3.**
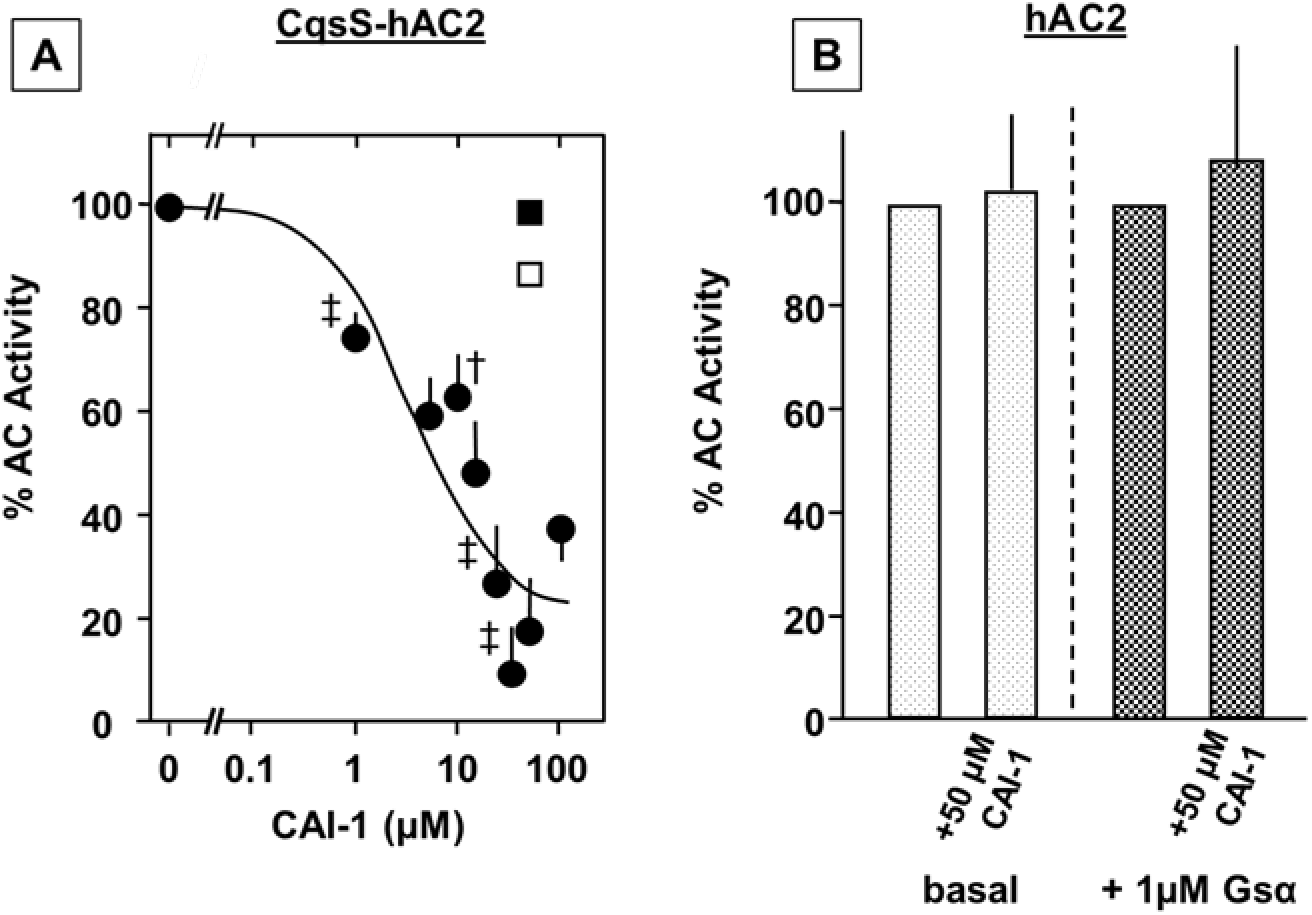
A) CAI-1 concentration-response curve for the inhibition of Gsα-stimulated CqsS-hAC2. 1 μM Gsα-stimulated activity above basal (100 %) was 33 ± 2.6 pmol cAMP·mg^−1^·min^−1^. The IC_50_ for CAI-1 was 6.3 μM. Error bars denote S.E.M. of 3-7 separate experiments (expressions). Significances: †: p <0.01; ‡: p < 0.001 compared to Gsα-stimulated activity (100%). For clarity, not all significances are indicated. **B) CAI-1 has no effect on Gsα stimulation of hAC2**. Basal (60 ± 20 pmol cAMP·mg^−1^·min^−1^) and 1 μM Gsα stimulated activities (2.7 ± 1.4 nmol cAMP·mg^−1^·min^−1^) were set at 100 %, respectively; CAI-1 was dissolved in DMSO, 2 % final DMSO in all assays. 5 independent assays from one Sf9 expression were carried out. Error bars denote SD.

### Gsα Stimulation of hAC2 is inhibited by human serum

The above data present a proof-of-concept experiment to demonstrate that a 2×6TM anchor domain can regulate formation of the catalytic dimer of hAC2. The data pose the questions: are mACs regulated in a similar manner and what are potential ligands in mammals and where to expect them? Considering that the mAC membrane anchors are isoform-specifically conserved, prospective ligands are predicted to be similarly primordial (Bassler et al., 2018, Ziegler et al., 2017). Because ligands supposedly access the mACs from the extracellular solvent space, they are expected to be systemically present in the extracellular fluid system of the body. Indeed, human serum significantly inhibited stimulation of hAC2 by 600 nM Gsα (and similarly did heat-inactivated human serum in which complement is inactivated) in a concentration-dependent manner (Fig. 4A). Serum albumin had no effect (Fig. 4A). Fetal bovine serum (FBS) was even more potent indicating a higher concentration of inhibitory factors and almost excluding immunoglobulins as potential ligands because the concentration of immunoglobulins in FBS is substantially lower compared to human serum (Fig. 4 A).

**Figure 4.**
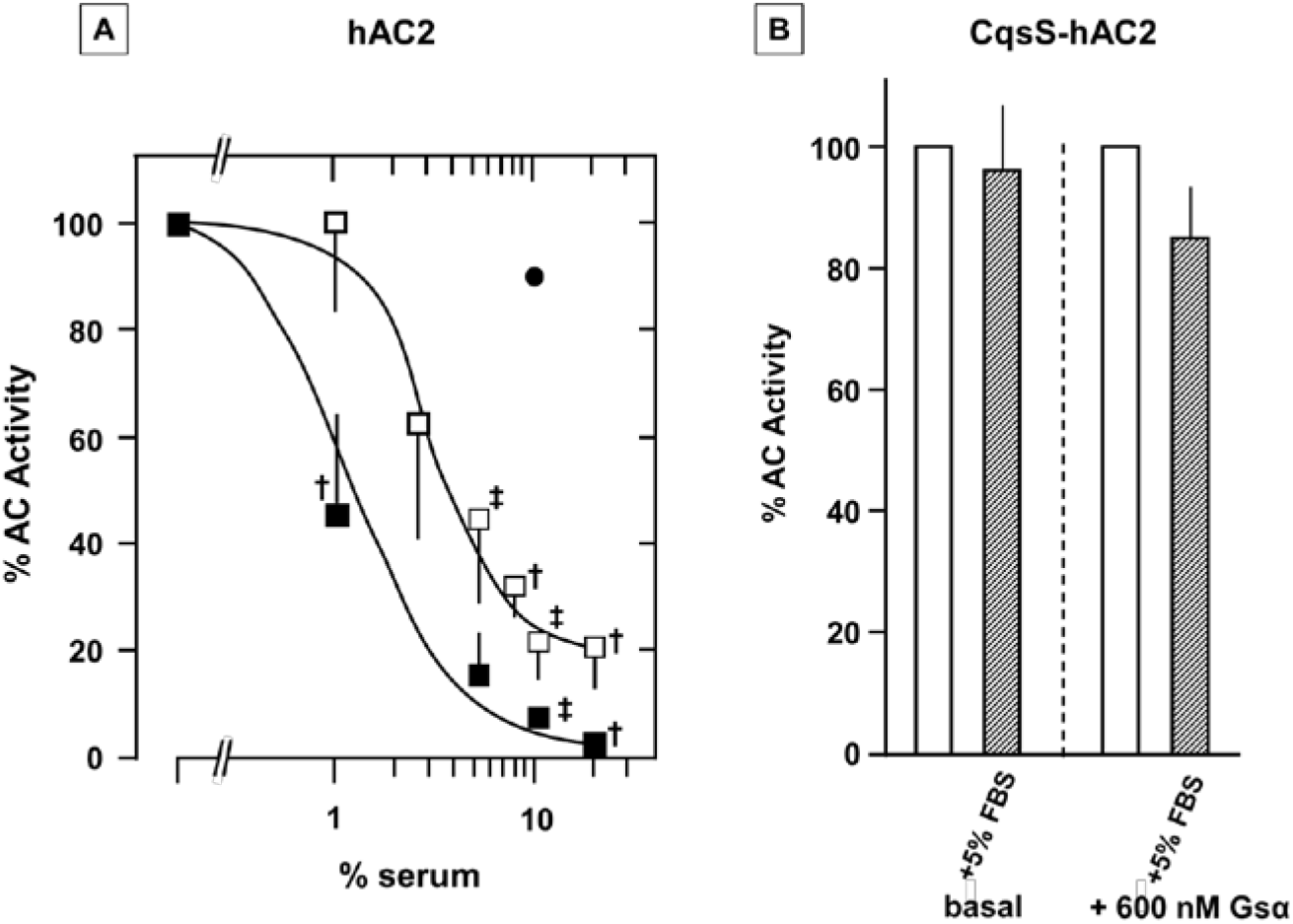
**A)** Inhibition of Gsα stimulation (600 nM) of hAC2 by human serum (□) and FBS (■); human serum albumin (●) at 45 μg/assay. Basal hAC2 activity was 30 pmol cAMP·mg^−1^·min^−1^, 100 *%* Gsα-stimulated activity was 1.6 nmol cAMP·mg^−1^·min^−1^. Error bars represent SD of 3-5 experiments. The difference between the serum concentrations for half maximal inhibition by human serum and FBS did not reach significance. **B)** The CqsS-hAC2 chimera was unaffected by 5% FBS in the assay (18.5 μg protein). Basal (open bars; 13.6 ± 2.2) and Gsα-stimulated activity (hatched bars; 27.2 ± 6.2 pmol cAMP·mg^−1^·min^−1^). Error bars represent SD of four assays. Significances: †: p<0.01 compared to respective 100 % control; ‡: p<0.001.

Although hAC2 activity with Mn^2+^-ATP was much higher than with Mg^2+^-ATP we preferred using Mg^2+^-ATP as the likely physiological divalent cation. Serum itself contains about 1.25 mM Mg^2+^ and 2.4 mM Ca^2+^, virtually excluding a chelating effect of divalent cations by serum addition and, in all likelihood, regulatory effects of Ca^2+^ (Csenker, Dioszeghy et al., 1982). The data suggested that potential receptor ligands for hAC2 are present in serum. We also carried out concentration-response curves for Gsα stimulation of hAC2 in the absence and presence of 5 and 10 % human serum (appendix fig. S1). In the presence of serum the efficacy of Gsα was substantially diminished, increasing the required Gsα concentrations for activation and diminishing the maximal responses (appendix fig. S1). This was similar to the results with the CqsS-hAC2 chimera (see Fig. 2A) suggesting similar mechanisms of action for CAI-1 and specific serum factors, respectively.

Stringent controls are required to unequivocally assign the effect of serum to the membrane anchors of hAC2: a) exclusion of a salt effect as serum contains around 120 mM NaCl, b) exclusion of an interference of serum components with the dimerization of the C1 and C2 catalytic domains, and c) exclusion of an interference in the interaction between the catalytic dimer and Gsα. 10 mM NaCl in the incubations, equivalent to 10% serum in the assays, did not impair hAC2 basal activity or Gsα stimulation. Similarly, using dialyzed serum did not abrogate inhibition of Gsα stimulation excluding interference by NaCl.

Basal activities of all mACs generally are rather low and difficult to determine reliably (Hildebrandt & Birnbaumer, 1983). Therefore, we increased the amount of hAC2 membrane protein 27-fold and examined the effect of serum. It inhibited basal hAC2 activity in a concentration-dependent manner with a half-maximal inhibition at 4.9 % serum whereas albumin did not (Appendix fig. S2).

This created a critical issue because components of human or fetal bovine serum might either interfere with dimerization of C1 and C2 domains or with activation of the dimer by Gsα. This was explored using CqsS-hAC2 in which the membrane anchors are from a *Vibrio* quorum-sensor whereas the cytosolic domains are from hAC2 (scheme in Fig. 1). In the CqsS-hAC2 chimera FBS did neither interfere with basal hAC2 activity nor with Gsα stimulation (Fig. 4B). Thus, we excluded an effect of serum on the catalytic hAC2 dimer or on Gsα activation of the dimer. The results demonstrated that the action of serum on basal hAC2 activity was contingent on the presence of the membrane anchor of hAC2.

Are the above results restricted to the hAC2 isoform or are they indicative of a more general regulatory mechanism applicable to all mACs? Based on pronounced sequence features the nine mACs are subclassified into four subclasses, AC 1, 3 and 8, AC 2, 4, and 7, AC 5 and 6, and the standalone isoform AC9 (Dessauer et al., 2017, Sunahara & Taussig, 2002, Willoughby & Cooper, 2007). We investigated hAC3, hAC5 and hAC9, i.e. one member of each subclass (including hAC2). Using appropriate enzyme concentrations, serum inhibited basal activities of hACs 3, 5, and 9 with hAC5 being less sensitive to inhibition compared to hAC3 and 9 (IC_50_ = 6.3 % serum) indicating either different concentrations of specific inhibitory factors or differences in affinity of a potentially common factor (Appendix fig. S3). Similarly, Gsα stimulation of hAC isoforms 3, 5, and 9 was inhibited by serum. The calculated IC_50_ concentrations ranged between 3 and 7 % (Fig. 5A). These insignificant differences were not surprising as the commercial human serum is mixed from adult human donors, certainly presenting various physiological states while donating blood.

**Figure 5.**
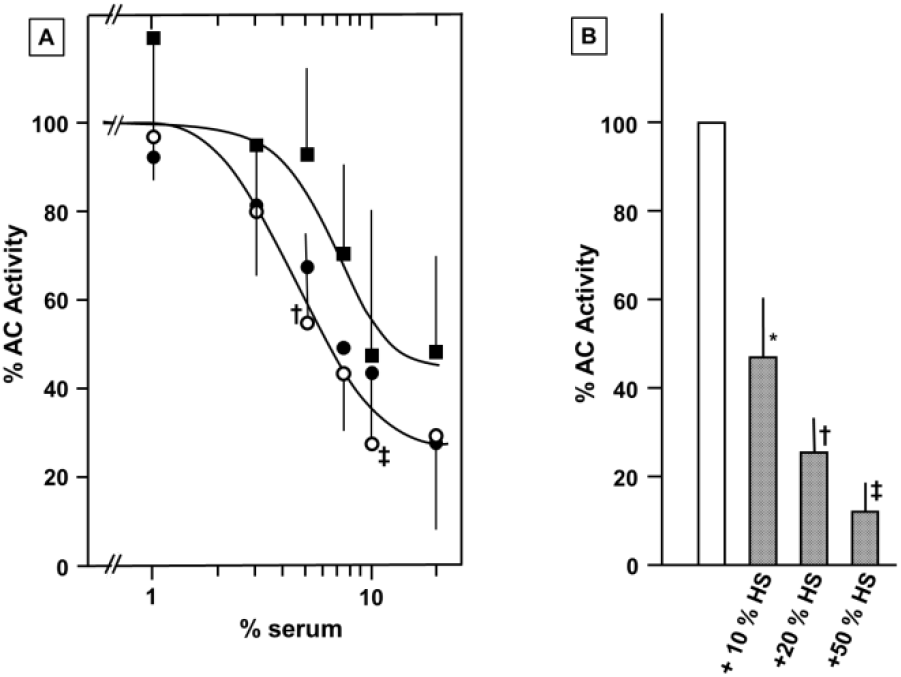
**A)** Serum inhibition of hACs 3, 5, and 9 stimulated by 600 nM Gsα (○: hAC 3; ●: hAC5; ■: hAC9). To depict the data in a single graph, activities of hACs 3, 5, and 9 stimulated by 600 nM Gsα were taken as 100% (hAC3 = 0.7, hAC5 = 4.5, AC9 = 1.5 nmol cAMP·mg^−1^·min^−1^; inhibition of basal AC activities see appendix fig. S3). **B)** Serum inhibition of AC activity in rat brain cortical membranes stimulated by 600 nM Gsα. 100% activity corresponds to 1.24 nmol cAMP·mg^−1^·min^−1^ (3.5-fold stimulation above basal; n=3). †: p<0.01: ‡: p<0.001 compared to 100 % control. Error bars denote SD of 3-4 experiments carried out with individual hAC expressions, respectively (for inhibition of basal activities see appendix fig. S4). For clarity, not all significances are indicated. 1% serum corresponds to addition of 51 μg protein).

Next, we investigated whether we might have dealt with a serum effect related to the heterologous expression of hACs in Sf9 cells. We prepared membranes from rat brain cortex which contain essentially all mAC isoforms (Sanabra & Mengod, 2011). Serum potently inhibited basal AC as well as Gsα stimulated activity, suggesting that the mACs in brain membranes are similarly regulated as individual AC isoforms expressed in Sf9 cells (Fig. 5B and appendix fig. S4).

mACs are known to be regulated by a number of cytosolic effectors such as Gβγ, calcium / calmodulin, or forskolin and several secondary modifications such as phosphorylation (Dessauer et al., 2017). These factors generally have divergent, isoform-specific effects, e.g. Gβγ is reported to enhance Gsα or forskolin-stimulated activities of mACs 2, 4, 5, 6, and 7, but to have no effect alone (reviewed in (Brand, Sadana et al., 2015, Willoughby & Cooper, 2007)). On the other hand, Gβγ inhibits mACs 1, 3, and 8, and is even reported to inhibit AC5 and 6 (Steiner, Avidor-Reiss et al., 2005, Tesmer & Sprang, 1998, Willoughby & Cooper, 2007). Similarly, calcium and calmodulin have isoform-specific inhibitory or activating effects (Willoughby & Cooper, 2007). In contrast, the plant diterpene forskolin uniformly activates mACs 1 to 8 and it potentiates Gsα activation (Dessauer et al., 2017). In crystal structures of the catalytic dimer, forskolin is bound within the catalytic cleft (Tesmer et al., 1997, Tesmer, Sunahara et al., 1999). Therefore, we examined whether serum affects the action of forskolin on hAC2, either alone or in conjunction with Gsα. 25 μM Forskolin stimulated hAC2 about 2.4-fold and human serum significantly inhibited activation (Fig. 6A). 25 μM Forskolin + 300 nM Gsα resulted in a 6.6-fold potentiation of activation of hAC2 and serum was significantly inhibitory as well (Fig. 6B).

**Figure 6.**
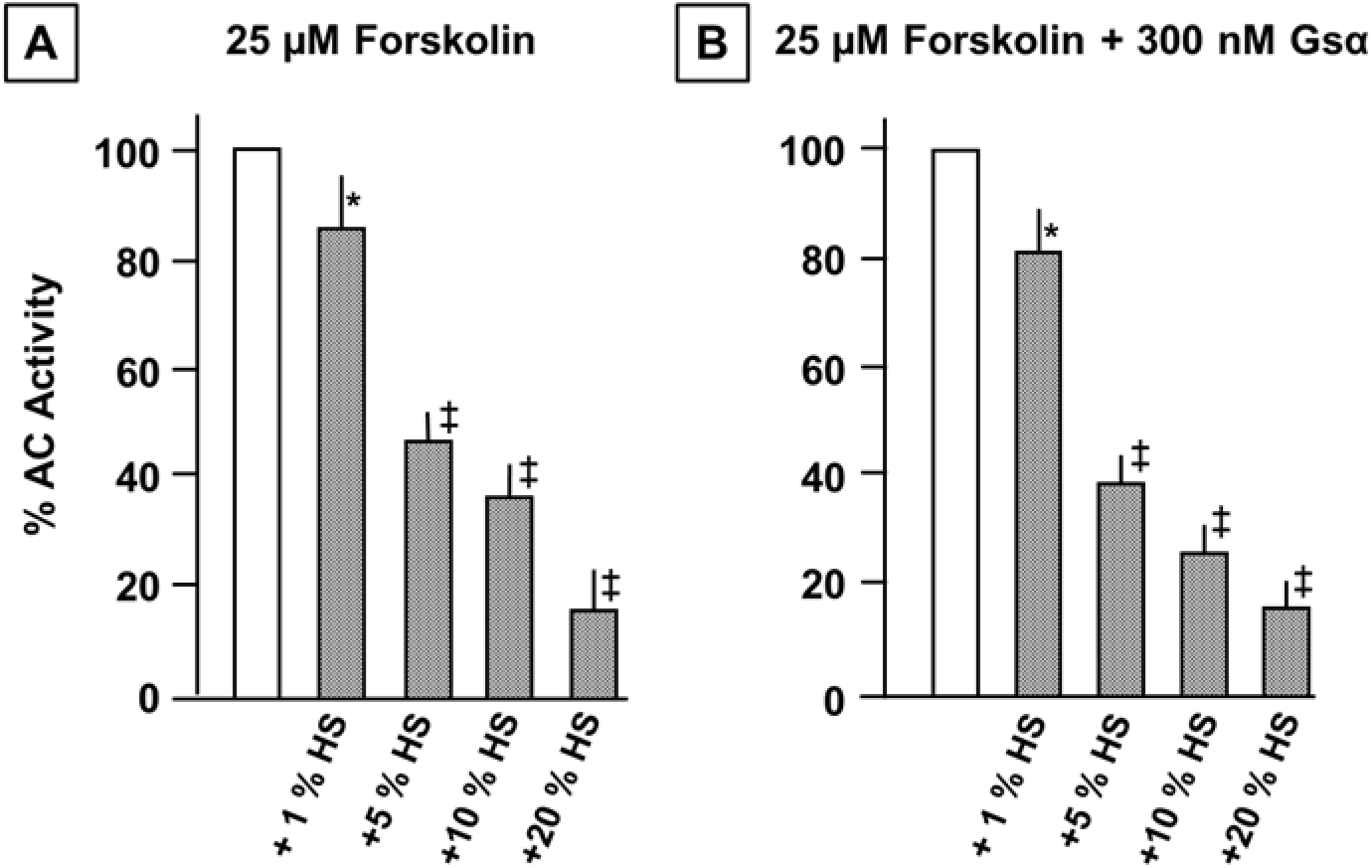
Serum inhibition of hAC2 activity stimulated by forskolin and forskolin + Gsα. **A)** Inhibition of 25 μM forskolin stimulated hAC2 activity (100 % corresponds to 0.19 nmol cAMP·mg^−1^·min^−1^, a 2.5-fold stimulation). **B)** Inhibition of hAC activity stimulated by 25 μM forskolin and 300 nM Gsα (100 % corresponds to 0.5 nmol cAMP·mg^−^ min^−^, a 6.6-fold stimulation). Error bars denote SD of four experiments with membranes from one Sf9 expression of hAC2. Significances: *: p<0.05; ‡: p<0.001.

Interestingly, also basal AC activity in rat cortical membranes was inhibited by serum in line with respective results of hACs 2, 3, 5, and 9 activities (Appendix fig. S2,3 and 4). This demonstrates that regulatory processes of mACs, which are mediated via direct effects on the cytosolic catalytic dimer, are affected by action of specific inhibitory factors present in serum acting via mAC membrane domains. The data support our suggestion that we were dealing with a novel general mechanism of mAC regulation.

## Discussion

Thus far, studies of regulation of mAC activity mostly dealt with regulation of the cytosolic catalytic dimer, primarily by uniformly activating Gsα and, secondarily, by variable other inputs (reviewed in (Dessauer et al., 2017, Sunahara & Taussig, 2002)). The biochemical data emanating from these studies are usually discussed by a two-state model, an active and an inactive state. In this respect, the 2×6TM anchors were considered inert. Potential roles for the membrane anchors were assigned to localization, e.g. in membrane rafts, or as potential interaction sites for scaffolding proteins (Crossthwaite, Seebacher et al., 2005, Li, Chen et al., 2012, Piggott, Bauman et al., 2008, Rich, Fagan et al., 2000). Our data demonstrate a new level of mAC regulation which is spatially distinct from the catalytic dimer, and, for the first time, confer a regulatory function to all mAC domains. In addition, the regulatory input via the AC membrane domains immediately suggests a possible explanation for the striking evolutionary conservation of the membrane anchors in an isoform-specific manner (Bassler et al., 2018) and requires expanding the previous two-state model of mAC regulation to a three-state model.

CqsS is a hexahelical quorum-sensing receptor from *Vibrio* sensing the extracellular ligand CAI-1 (Ng, Perez et al., 2011, Ng, Wei et al., 2010). It is isosteric to a 6TM domain of the pseudoheterodimeric mACs (Beltz et al., 2016). In the CqsS-hAC2 chimera, we observed that CAI-1 attenuated Gsα stimulation in an unequivocally receptor-mediated process (Fig. 2). CAI-1 had no such effect on Gsα-stimulated activity of the hAC2 holoenzyme, because the hAC2 membrane anchor lacks the functionality to sense CAI-1 (Fig. 2, 3). However, stimulation of hAC2 by Gsα was inhibited by serum (Fig. 4A). The effect was dependent on the membrane domain from hAC2 because serum did not affect Gsα stimulation of CqsS-hAC2 as the CqsS receptor cannot sense signaling components present in mammalian serum (Fig. 4B). In addition, this demonstrated that serum did not affect dimerization of C1 and C2 (Fig. 4B). Serum albumin, the major protein in serum, had no effect suggesting the presence of specific, as yet unidentified inhibitory components in serum (Fig. 4A). Inhibition of Gsα stimulation by serum was further demonstrated for hAC isoforms 3, 5, and 9, thus covering one isoform from each mAC subclass (Fig. 5A). Likewise, serum inhibited basal and Gsα stimulated mAC activity present in rat brain cortical membranes (Fig. 5B and appendix fig. S4), virtually excluding the possibility of an artifact due to heterologous expression of hACs in Sf9 insect cells. Conceptually, a regulatory input from the extracellular space should affect all cytosolic regulatory inputs impinging upon the catalytic dimer. This was verified with forskolin which stimulates cytosolic catalytic dimer (exception mAC9). Forskolin activation of hAC2 ± Gsα was inhibited by serum (Fig. 6B).

### A three-state model of adenylate cyclase regulation

Based on these data and equilibrium thermodynamic considerations we propose a novel formal concept of regulation of mACs which encompasses all available biochemical, pharmacological and structural data (Fig. 7). Three distinct basal states of mACs exist in equilibrium, state A (inactive), state B (inactive), and state C (active). States A and B differ in the conformational flexibility of their catalytic C1/C2 domains. In state A, the catalytic domains are conformationally constrained and cannot form an active dimer. In state B, the catalytic domains are conformationally unconstrained, yet because of their low affinity for each other, they only occasionally collapse into an active dimer (state C). The highly transient state ‘C’ is responsible for the very low basal activity observed in all mACs.

**Figure 7.**
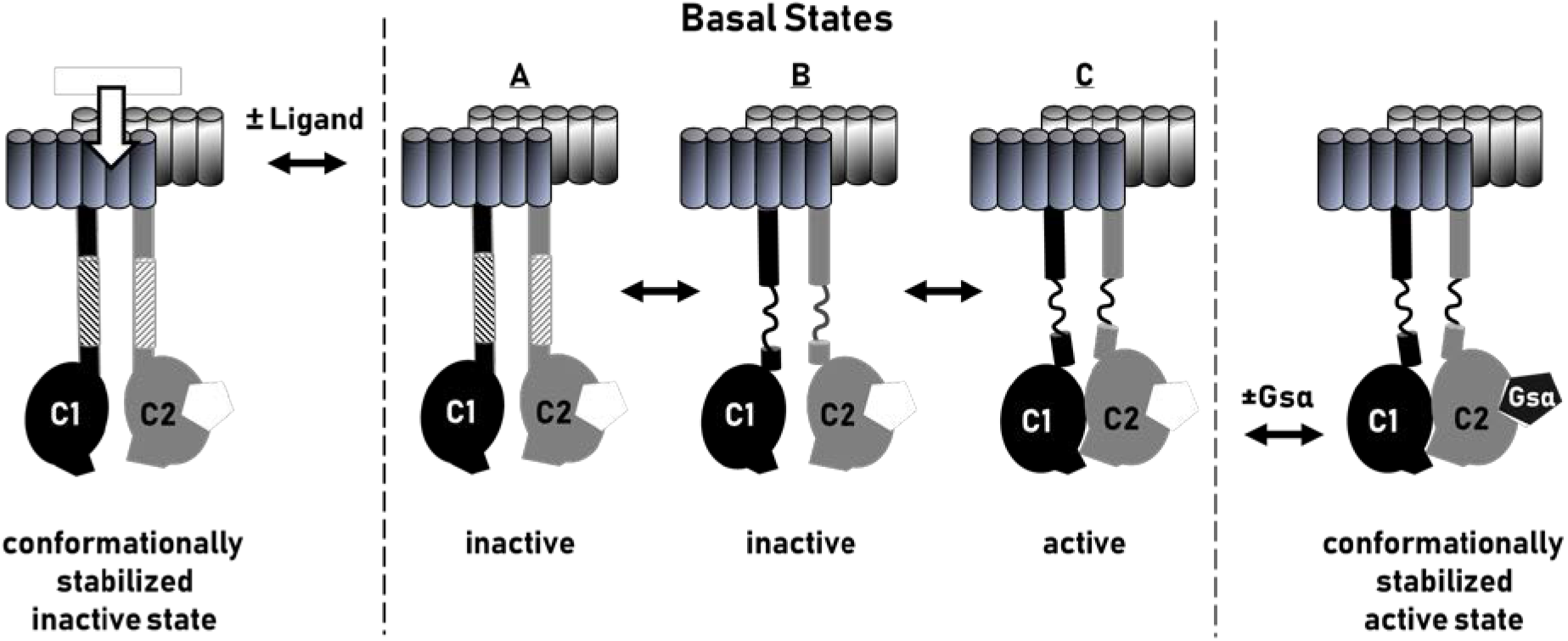
Scheme of regulation of mammalian adenylate cyclases. Three basal states are in a thermodynamic equilibrium, two inactive, A and B, and one active state, C. The constant low basal enzyme activity is due to fractional formation of an active dimer as symbolized in C. Contact with Gsα in the cytosol stabilizes the active ‘C’-state (conformationally stabilized active state, at far right). Conversely, binding of as yet unknown ligands at the extracellular side of the membrane anchor stabilizes the inactive state A (conformationally stabilized inactive state; at far left).

Constraining structural flexibility by binding of a ligand at the extracellular side shifts the equilibrium to the inactive ‘A’-state (Fig. 7, far left) attenuating basal AC as well as Gsα stimulated activities in hAC2, 3, 5, 9, and in rat brain cortical membranes (Fig. 4, 5 and suppl. mat. fig. 2–4). Forskolin reversibly increases the apparent affinity of C1 and C2 about tenfold, i. e. it stabilizes the ‘C’-state. The ‘C’-state is further stabilized by binding of Gsα at the cytosolic dimer thus activating mACs and potentially acting in concert with forskolin (Fig. 7, far right).

The proposed three-state model suggests the existence of an allosteric linkage in mACs, in which the membrane anchors, as receptors, transduce extracellular signals across the cell membrane to the cytosolic catalytic dimer. This way, each mAC isoform can be addressed individually by an extracellular ligand and primed for a physiologically measured GPCR/Gsα response. Such a regulatory network would explain the conundrum of why often multiple Gsα-stimulated mAC isoforms are expressed in a single cell. The model is in agreement with the recently published structure of a bovine AC9 isoform which contains features compatible with signal transduction between membrane anchor and catalytic dimer (Qi et al., 2019). We are aware of the fact that the equilibrium of states will be subject to ambient conditions such as ion and substrate concentrations, membrane charge and membrane potential. Thus, the model is open to modifications without losing its conceptual validity. Similarly, the lack of chemically identified ligands neither impaired establishing nor does it affect the validity of the three-state model. Considering the isoform-specific conservation of AC membrane anchors nine ligands are expected. These ligands in conjunction with the respective receptors will define a regulatory system which affects GPCR/Gsα actions executed via mACs. In summary our study not only unequivocally designates AC membrane anchors as orphan receptor but also it identifies the pathway to take for identification of the ligands

### Materials and methods

CqsS of *V. harveyi* facc. # AAT86007) and hAC2 and hAC9 (acc. # Q08462 and NM_001116.3) sequences were used. Position Phe166 in CqsS has been shown to be critical for quorum sensing and was mutated to Leu (F166L) according to Beltz et al. (Beltz et al., 2016, Ziegler et al., 2017). Radiochemicals were from Hartmann Analytic and Perkin Elmer. Enzymes were from either New England Biolabs or Roche Molecular. Other chemicals were from Sigma, Merck and Roth. CAI-1 was synthesized in-house according to (Ng, Perez et al., 2012). The cAMP assay kit is from Cisbio (Berlin). The constitutively active GsαQ227L point mutant was a gift from Dr. C. Kleuss, Berlin (Diel, Klass et al., 2006). It was expressed in *E. coli* and purified by affinity chromatography via its N-terminal His_6_-tag and by size-exclusion chromatography as described. Human serum albumin (catalog # A3782) was from Sigma-Aldrich (Taufkirchen, Germany), fetal bovine serum was from Gibco, Life Technologies, Darmstadt, Germany (catalog #: 10270; lot number: 42Q8269K).lot number: 1908933). Plasmid construct Several CqsS_hAC2-C1_CqsS_hAC2-C2 chimeras were generated using standard molecular biology methods. DNA fragments and vectors were restricted at their 5’-BamHI or EcoRI and 3'-HindIQ sites and inserted into pQE80L (Δ Xhol; Δ Ncol). Gly-Ser and Arg-Ser were introduced to generate required internal restriction sites. All constructs carried N-terminal MRGS-hexa-His-tags to examine potential proteolysis by Western blotting. The fidelity of all constructs was confirmed by double-stranded DNA sequencing. The construct used in this study was: MRGSHis6-GS–CqsS (F166L)1-181-hAC2221-603-RS-CqsS (F166L)1-181-hAC2836-1091. For virus production, hAC2 was inserted into pLIB. The plasmid was amplified in *E. coli* XL1blue and transformed into *E. coli* EMBacY cells generating the bacmid for Sf9 transfection. Genes for all hACs were bought from GenScript.

### Plasmid construct

CqsS-hAC2 was generated using standard methods. 5’-BamHI or EcoRI and 3-HindIII sites restriction sides were used and inserted into pQE80_L_ (Δ XhoI; Δ NcoI). Gly-Ser and Arg-Ser were introduced for internal restriction sites. An N-terminal MRGS-hexa-His-tag was used to for Western blotting to analyze potential proteolysis. It was absent. The fidelity of the construct was confirmed by double-stranded DNA sequencing. The construct boundaries were: MRGSHis6-GS—CqsS-(F166L)_1-181_-hAC222_1-603_-RS-CqsS-(F166L)_1-181_-hAC2_836-109_. Genes for hACs 2, 3, 5, and 9 were obtained from GenScript. For virus production, hAC genes were inserted into pLIB. The plasmid was amplified in *E. coli* XL1blue and transformed into *E. coli* EMBacY cells generating the respective bacmids for Sf9 transfection.

### Protein expression

CqsS-hAC2 was transformed into *E. coli* BL21 (DE3). Strains were grown overnight in Luria-Bertani broth (LB medium) at 30°C with 100 μg/ml ampicillin. 200 ml LB medium or M9 with glycerol replacing glucose were inoculated with a preculture (to A600 of 0.2) and grown at 30-37°C in the presence of antibiotics. At an A600 of 0.9 (for M9 medium) / 0.7 (for LB medium), the temperature was lowered to 22°C and expression was initiated by addition of 500 μM isopropyl β-D-1-thiogalactopyranoside (IPTG) for 4 hrs. Cells were harvested by centrifugation, washed once with 50 mM Tris/HCl, 1mM EDTA, pH 8, and stored at −80°C. For preparation of cell membranes, cells were suspended in lysis buffer (50 mM Tris/HCl, 17 0.021 % thioglycerol, 50 mM NaCl, pH 8) containing complete protease inhibitor (in 50ml) cocktail (Roche Molecular) and disintegrated with a French press (1100 psi). After removal of cell debris (4.300 x g, 30min, 4°C) membranes were collected at 100000 x g (1h at 4°C). Membranes were suspended in buffer (40 mM Tris/HCl, 0.016 % thioglycerol, 20 % glycerol, pH8) and assayed for AC activity. Multiple expressions were carried out. Virus-infected Sf9 cells expressing hAC2, 3, 5, and 9 were grown in Sf900 III medium which contains no ingredients derived from animal sources, harvested after three days and membranes were isolated, stored at −80 °C and used for testing. hAC isoforms from single Sf9 transfections were used, respectively.

### Adenylate cyclase assay

Activity of CqsS-hAC2 was routinely assayed for 15 min at 30°C-37°C in 100μl with 40 μg membrane protein, 50 mM Tris/HCl pH 8.3, 5 mM MnCl_2_, 6 mM creatine phosphate, 230 μg/ml creatine kinase, 750 μM [α-^32^P]-ATP, and 2 mM [2,8-^3^H]-cAMP to monitor yield during cAMP purification (Salomon, Londos et al., 1974). With Mg^2+^ as cation basal and stimulated CqsS-hAC2 AC activities were lower. Substrate conversion was kept below 10%. CAI-1 was dissolved in DMSO. Incubations with DMSO were carried out as controls.

Activity of hACs was determined in a volume of 10 μl using 1 mM ATP, 2 mM MgCl_2_, 3 mM creatine phosphate, 60 μg/ml creatine kinase, 50 mM MOPS, pH 7.5 using an Assist-Plus pipetting robot (Integra Biosciences, Germany) and a cAMP assay kit from Cisbio (Codolet, France) according to the supplier’s instructions (see controls of standard curves in appendix fig. S5).

### Western blot analysis

For Western blotting an RGS-His_4_-antibody (Qiagen) and a 1:2500 dilution of the fluorophore-conjugated secondary antibody Cy3 (ECL Plex goat-α-mouse IgG-Cy3, GE Healthcare) was used. Proteolysis was not observed.

## Data analysis and statistical analysis

All incubations were in duplicates (CqsS-hAC2) or triplicates (hACs). S.E.M values are given for experiments with CqsS and refer to separately expressed and analyzed membrane proteins. S.D. values apply hACs expressed once in Sf9 insect cells the membranes of which were used throughout. Student t-test is use for analysis. Data analysis was with GraphPad prism 8.1.2.

## Author contributions

AS, MF and JG designed, carried out and analyzed experiments; JES, concept, analyzed data and wrote manuscript together with all others.

## Appendix

**Figure S1.**
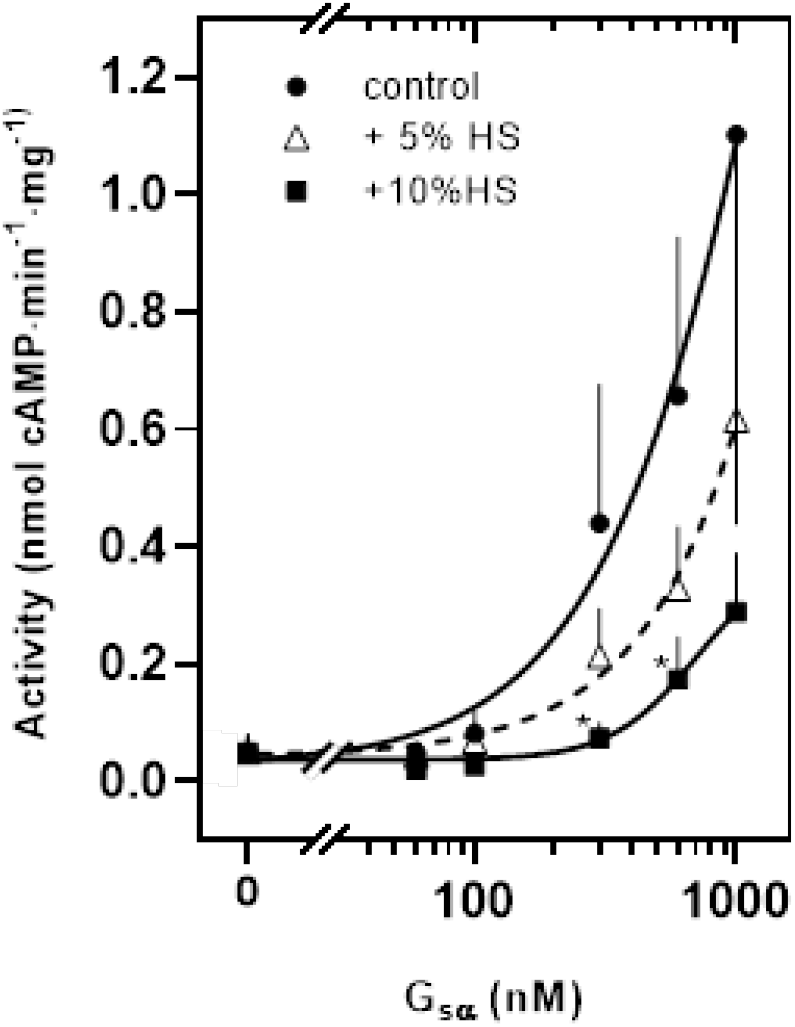
Gsα concentration-response curve of hAC2 in presence of 5 and 10 % human serum, respectively. Significances: *: p < 0.05 compared to control. Error bars denote SD of 3 experiments.

**Figure S2.**
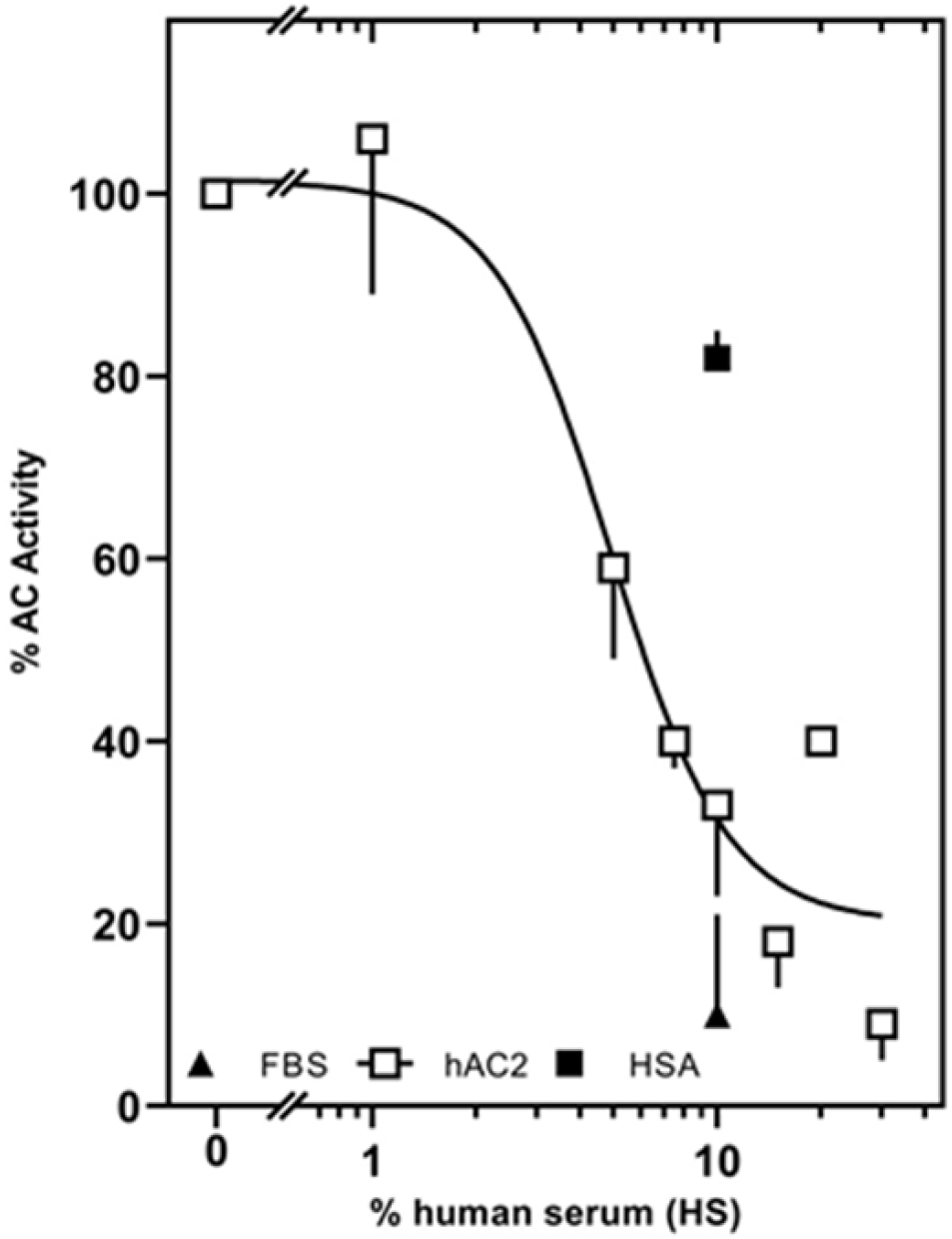
Inhibition of basal activity of hAC2 by human serum. Basal activity was 0.038 ± 0.006 nmol cAMP·mg^−1^·min^−1^.IC_50_ concentration determined by graph-pad was: 4,9% HS. 2-5 Experiments were carried out, error bars denote SD

**Figure S3.**
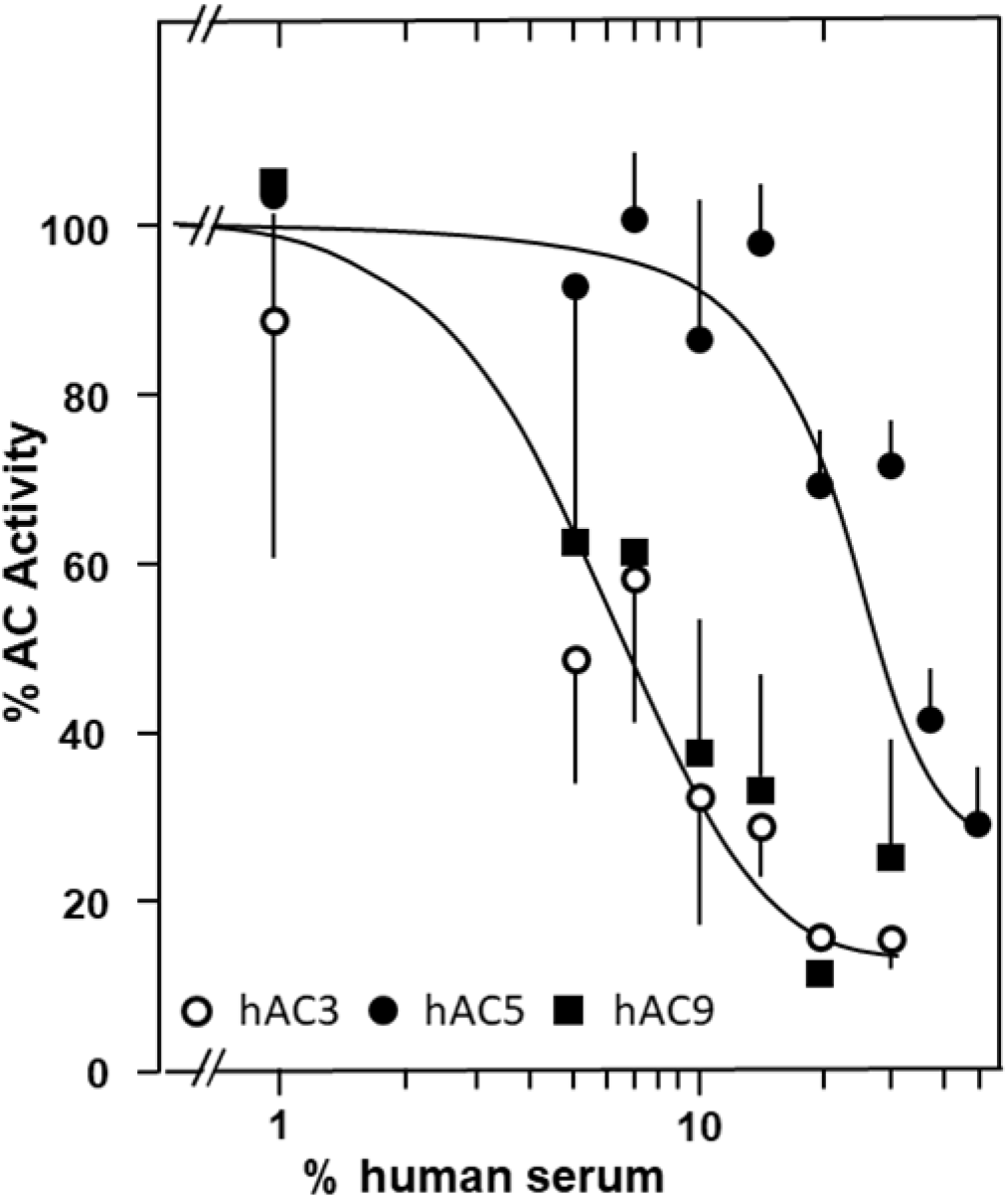
Inhibition of basal activity of hACs 3, 5, and 9 by human serum. 2-5 Experiments were carried out, Error bars denote SD. Basal activities were: hAC3 0.025 ± 0,007; hAC5 0.049 ± 0.013; hAC9 0.150 ± 0.047 nmol cAMP·mg^−1^min^−1^ IC_50_ concentrations determined by graph-pad were: 6.0%, about 40% and 5.9%, respectively

**Figure S4.**
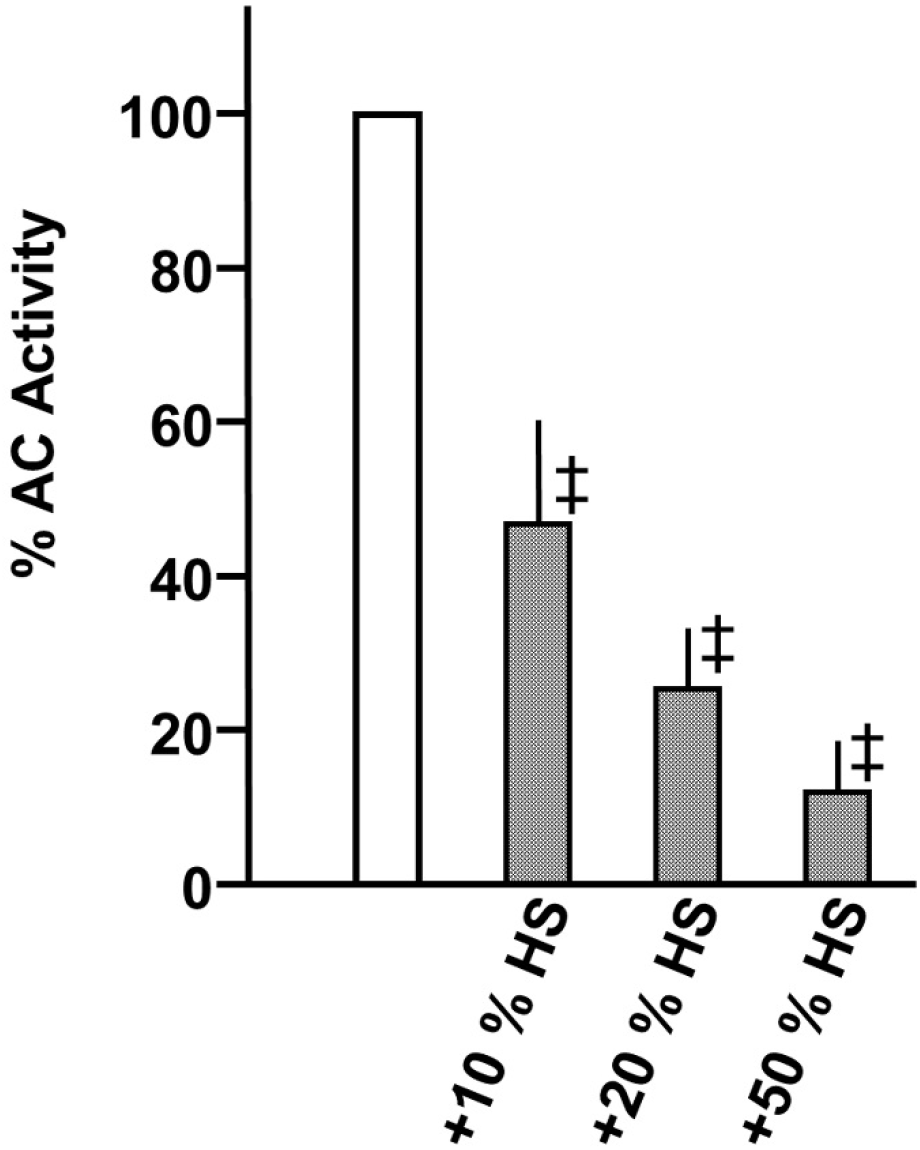
Inhibition of basal AC activity in a rat brain cortical membrane preparation. Basal activity (100 %) was 0.33 nmol cAMP·mg^−1^min^−1^. n = 3, Error bars denote SD. Significance: ‡: p< 0.001

**Figure S5.**
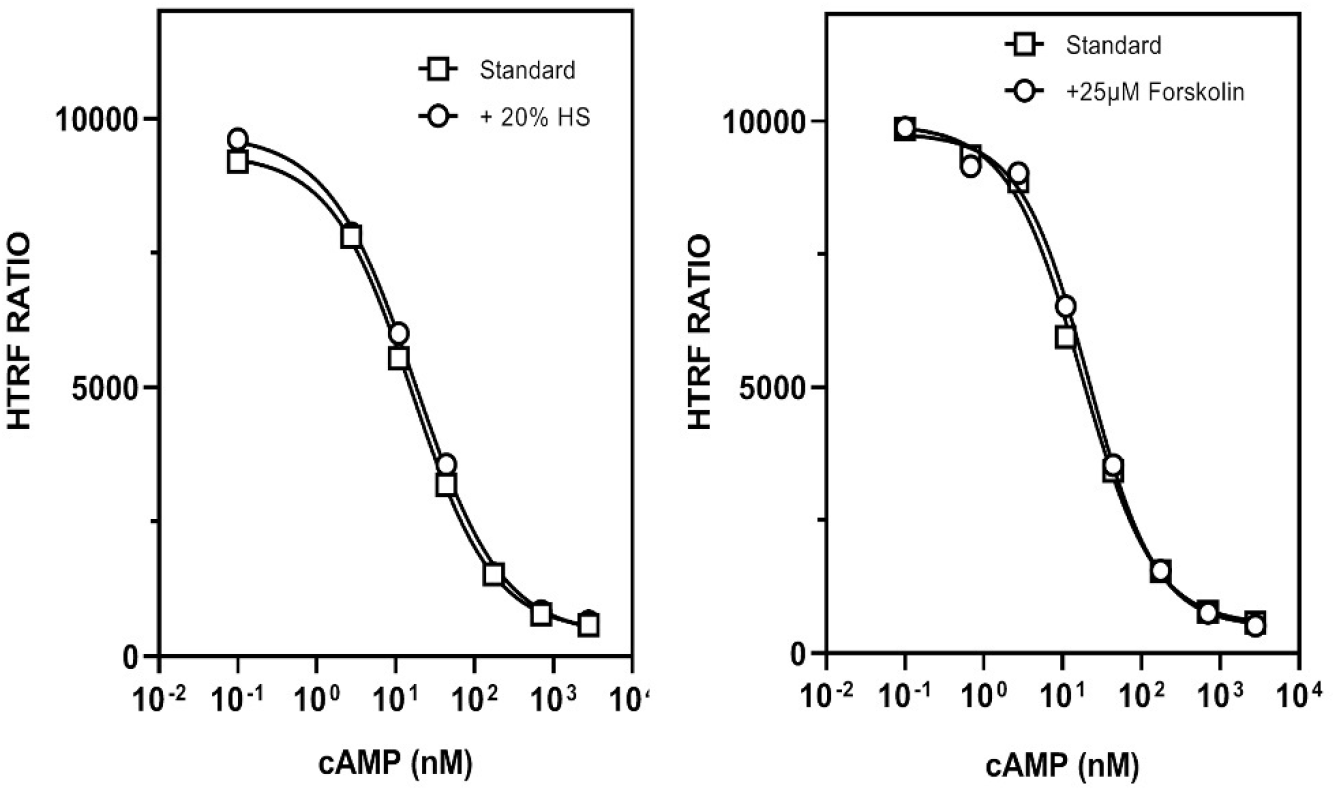
Standard curves for cAMP determination generated with the homogenous-time-resolved fluorescence assay (HTRF) from Cisbio in presence of 20 % human serum or 25 μM forskolin. No interference of serum and forskolin (and other agents used in respective AC assays such as Gsα at 1 μM or DMSO at 2%) was observed.

